# Zinc Import Mediated by AdcABC is Critical for Colonization of the Dental Biofilm by *Streptococcus mutans* in an Animal Model

**DOI:** 10.1101/2021.01.22.427828

**Authors:** Tridib Ganguly, Alexandra M. Peterson, Jessica K. Kajfasz, Jacqueline Abranches, José A. Lemos

## Abstract

Trace metals are essential to all domains of life but toxic when found at high concentrations. While the importance of iron in host-pathogen interactions is firmly established, contemporary studies indicate that other trace metals, including manganese and zinc, are also critical to the infectious process. In this study, we sought to identify and characterize the zinc uptake system(s) of *S. mutans*, a keystone pathogen in dental caries and a causative agent of bacterial endocarditis. Different than other pathogenic bacteria, including several streptococci, that encode multiple zinc import systems, bioinformatic analysis indicated that the *S. mutans* core genome encodes a single, highly conserved, zinc importer commonly known as AdcABC. Inactivation of the genes coding for the metal-binding AdcA (Δ*adcA*) or both AdcC ATPase and AdcB permease (Δ*adcCB*) severely impaired the ability of *S. mutans* to grow under zinc-depleted conditions. Intracellular metal quantifications revealed that both mutants accumulated less zinc when grown in the presence of a sub-inhibitory concentration of a zinc-specific chelator. Notably, the Δ*adcCB* strain displayed a severe colonization defect in a rat oral infection model. Both Δ*adc* strains were hypersensitive to high concentrations of manganese, showed reduced peroxide tolerance, and formed less biofilm in sucrose-containing media when cultivated in the presence of the lowest amount of zinc that support their growth, but not when zinc was supplied in excess. Collectively, this study identifies AdcABC as the lone high affinity zinc importer of *S. mutans* and provides preliminary evidence that zinc is a growth-limiting factor within the dental biofilm.

## Introduction

The first-row *d*-block elements iron, manganese and zinc are essential to all forms of life by serving structural, catalytic and regulatory functions to numerous biological processes. However, when in excess, these biometals are toxic such that their cellular flux and allocation must be tightly regulated. This Goldilocks paradox creates an opportunity for vertebrate hosts to deploy either metal sequestration or metal intoxication strategies to combat microbial infection; an active process known as nutritional immunity (Kehl-Fie & Skaar, 2010). Nutritional immunity strategies include mobilization of metal-chelating proteins to infected tissues such as iron-sequestering transferrin and lactoferrin, and neutrophil-secreted calprotectin which is responsible for restricting manganese, zinc and, in certain environments, iron availability (Nakashige, Zhang, Krebs, & Nolan, 2015; Zackular, Chazin, & Skaar, 2015). Paradoxically, the host can increase cytosolic zinc or mobilize copper or zinc into the phagosolysosome creating a toxic environment for intracellular bacteria (Sheldon & Skaar, 2019). To overcome metal limitation during infection, bacteria rely on the expression of surface-associated metal importers with some organisms also secreting small metal-binding molecules (metallophores) that tightly bind trace metals that are then reinternalized via specific transporters (Palmer & Skaar, 2016). Importantly, *in vivo* studies have shown that metal import systems are critical to bacterial virulence (Fischer et al., 2016; Garcia, Brumbaugh, & Mobley, 2011; Koh et al., 2015; Mastropasqua et al., 2017; Ong, Berking, Walker, & McEwan, 2018).

While the central role of iron in host-pathogen interactions has been known for decades, the importance of manganese and zinc homeostasis to bacterial pathogenesis is a relatively newer development (Kehl-Fie & Skaar, 2010; Lonergan & Skaar, 2019). In bacteria, manganese is the co-factor of enzymes involved in DNA replication, central metabolism and critical to activation of oxidative stress responses of gram-positive bacteria (Juttukonda & Skaar, 2015). Zinc is the second most abundant metal co-factor and is incorporated into 5-6% of all bacterial proteins, including central metabolism enzymes, regulatory proteins and metalloproteases (Lonergan & Skaar, 2019; Rahman & Karim, 2018). In addition, zinc plays an additional role in host-pathogen interactions by stimulating innate and adaptive immune cell function and, as indicated above, through mobilization within phagosomes to intoxicate invading pathogens (Lonergan & Skaar, 2019; Sheldon & Skaar, 2019; Subramanian Vignesh & Deepe, 2016). Different than iron and manganese, zinc does not undergo redox-cycling, but can form tighter and stabler interactions with metal ligands. As a result, zinc can occupy metal-binding residues of non-cognate metalloproteins inhibiting or reducing their activities; a process known as protein mismetallation (Chandrangsu & Helmann, 2016; Imlay, 2014).

A resident of dental plaque, *Streptococcus mutans* is a keystone pathogen in dental caries due to its ability to modify the oral biofilm architecture and environment in a way that facilitates the proliferation of acidogenic and aciduric bacteria at the expense of health-associated bacteria (Bowen, Burne, Wu, & Koo, 2018; Lemos et al., 2019). In addition to dental caries, *S. mutans* is a causative agent of infective endocarditis, a life-threatening infection of the endocardium (Pant et al., 2015). Once established in the oral biofilm, *S. mutans* metabolizes dietary carbohydrates, in particular sucrose, to produce an acidic biofilm matrix that is conducive to the growth of other cariogenic organisms (Lemos et al., 2019). Despite the importance of manganese and zinc to bacterial physiology, the mechanisms utilized by oral bacteria to maintain manganese and zinc homeostasis and their significance to polymicrobial and host-pathogen interactions in the oral cavity are poorly understood. Recently, we characterized the major manganese transporters of *S. mutans* and showed that maintenance of manganese homeostasis is critical for the expression of major virulence attributes of *S. mutans,* including the ability to tolerate acid and oxidative stresses and to form biofilms in a sucrose-dependent manner (Kajfasz et al., 2020). In this study, we sought to identify and characterize the zinc uptake system(s) of *S. mutans* and then determine the significance of zinc acquisition to the pathophysiology of *S. mutans*.

## Results

### *AdcABC is the main transporter responsible for zinc uptake in* S. mutans

In other streptococci, zinc acquisition is mediated by an ABC-type transporter known as AdcABC whereby AdcA is the zinc-binding lipoprotein, AdcB a membrane permease and AdcC a cytoplasmic ATPase (Bayle et al., 2011; Burcham et al., 2020; Loo, Mitrakul, Voss, Hughes, & Ganeshkumar, 2003; Ong et al., 2018). The *adcABC* genes are regulated by *adcR*, a metalloregulator from the MarR family, which is located immediately upstream and, depending on the streptococcal species, co-transcribed with *adcCBA* or *adcCB* (Fig. 1) (Reyes-Caballero et al., 2010). Through BLAST search analysis, we identified homologues of the *adcR*, *adcA*, *adcB* and *adcC* genes in the core genome of *S. mutans*. Similar to *S. agalactiae* and *S. pyogenes*, the gene coding for the zinc-binding AdcA lipoprotein is not co-localized with *adcR, adcB* and *adcC* being located as a monocistronic transcriptional unit elsewhere in the chromosome (Fig. 1). In addition, while the majority of streptococcal genomes encode two copies of *adcA*, dubbed *adcA* and *adcAII* (Bayle et al., 2011; Bersch et al., 2013; Brown et al., 2016; Burcham et al., 2020; Loisel et al., 2011; Loisel et al., 2008; Moulin et al., 2016), the genome of *S. mutans* encodes a single *adcA* gene copy, which is similar in size and more closely-related to the *adcA* gene genetically-linked to *adcBC* than to *adcAII* (Fig. 1). In most cases, the streptococcal *adcAII* is genetically coupled to metal-binding poly-histidine triad (*pht*) genes, which encode proteins that are thought to scavenge zinc outside the cell shuttling it to either AdcAII and, possibly, AdcA for internalization (Bersch et al., 2013; Loisel et al., 2011; Loisel et al., 2008; Zheng et al., 2011). While most streptococci encode at least one *pht* gene copy, bioinformatic analysis indicate that the core *S. mutans* genome lacks *pht* homologs. Similar to other streptococcal AdcA, the *S. mutans* AdcA has two zinc-binding domains that are characteristic of the genus: an N-terminal tertiary scaffold consisting of 3 histidine and 1 glutamic acid residue (H71, H149, H212, and E298) and a C-terminal ZinT domain consisting of three histidine residues (H461, H470 and H472) (Fig. 2 and S1). In addition, there are three additional conserved histidine residues (H71, H149 and H213) that aid in metal recruitment by AdcA (Cao et al., 2018).

**Figure 1.**
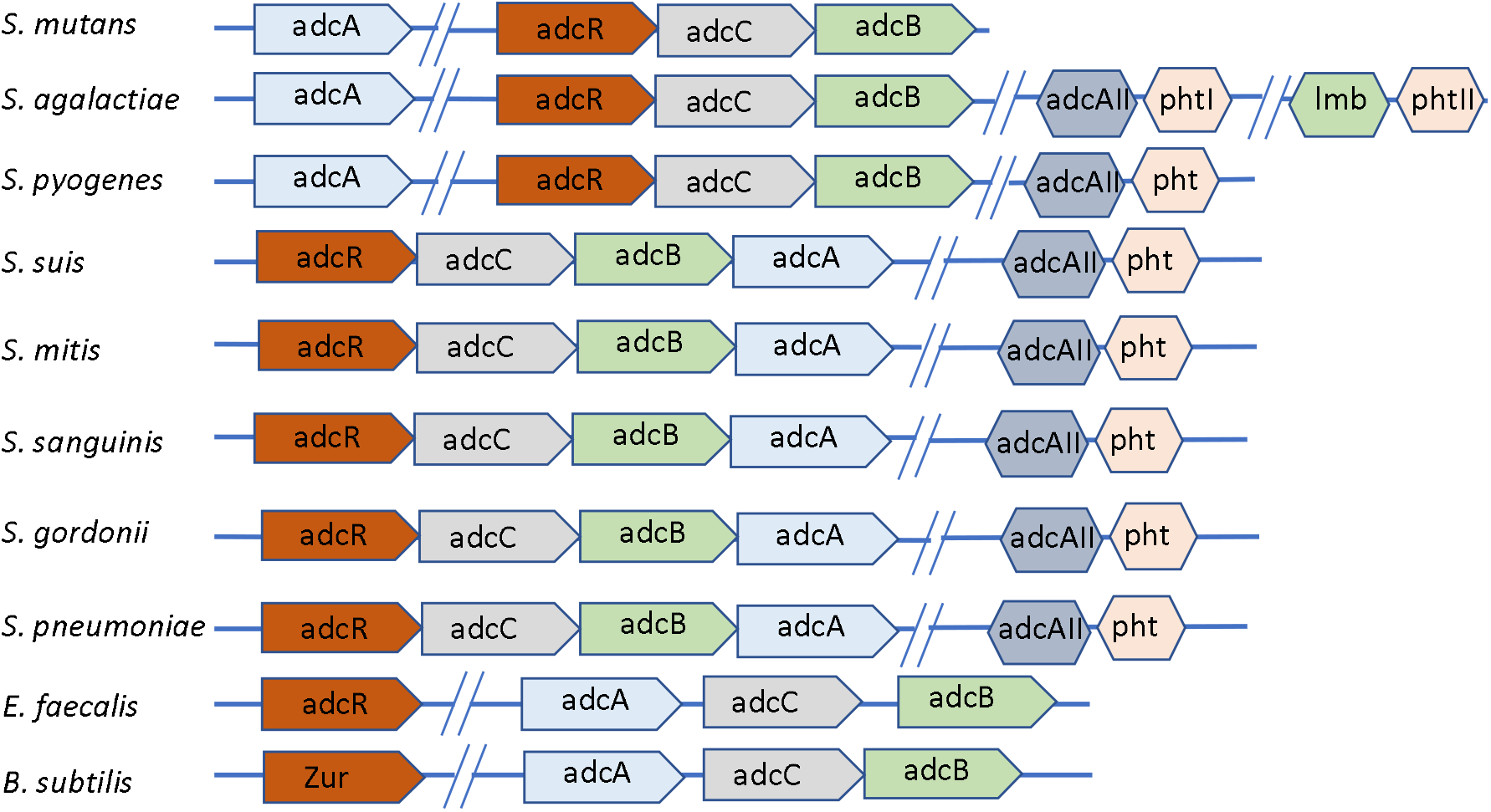
Genetic organization of the *adcABC* genes and other known zinc import systems in selected gram-positive bacteria.

**Figure 2.**
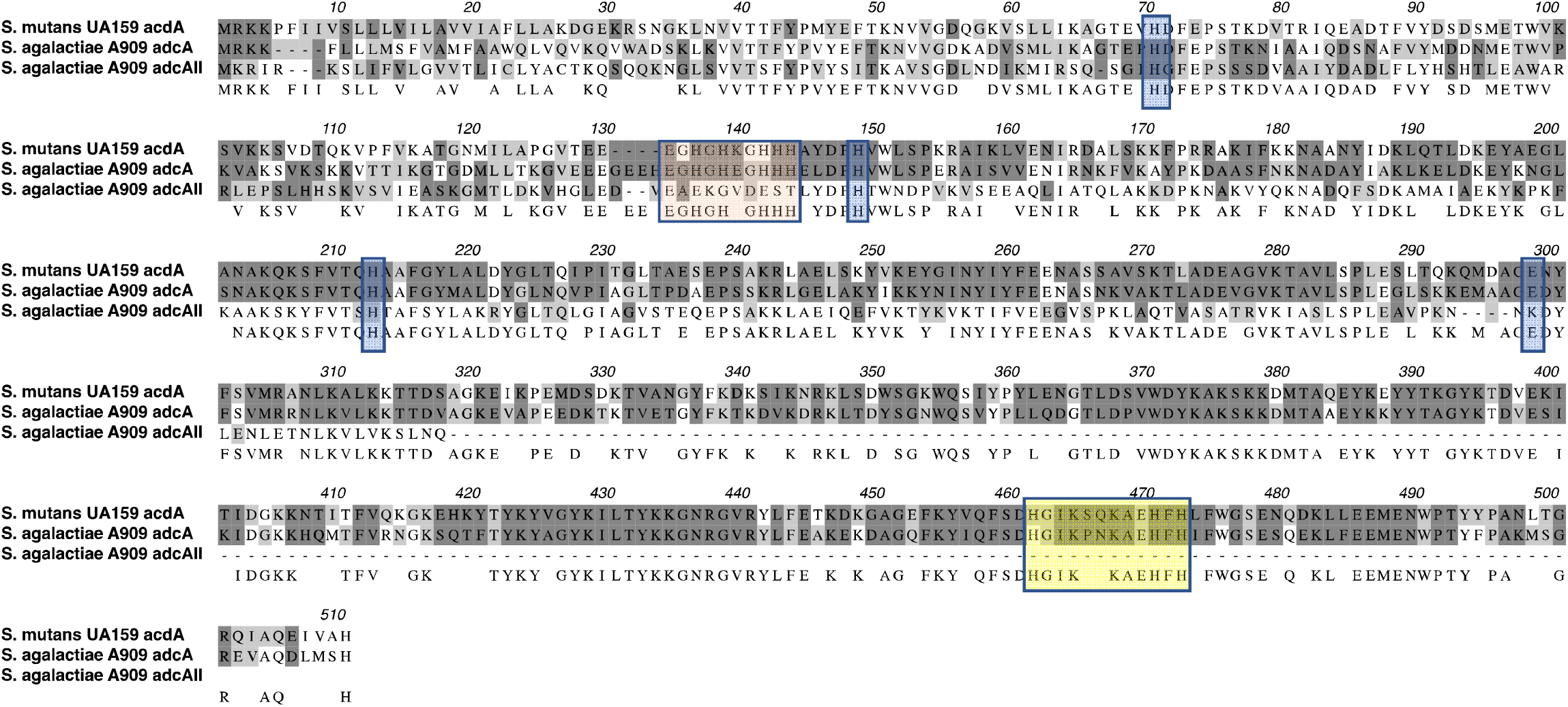
Alignment of AdcA of *S. mutans* with AdcA and AdcAII proteins of *S. agalactiae*. Dark and light grey shades represent identical and similar residues, respectively. Orange shaded residues are the N-terminal histidine rich metal-binding motif, yellow boxed residues depict the C-terminal ZinT domain while the glutamic acid and additional histidine residues that aid in metal recruitment are indicated in blue shades.

To evaluate the significance of zinc acquisition to the pathophysiology of *S. mutans*, we generated strains lacking *adcA* (Δ*adcA*) or both *adcC* and *adcB* (Δ*adcBC*) in the parent strain UA159 and then tested their ability to grow under zinc-replete or zinc-depleted conditions. Both Δ*adcA* and Δ*adcBC* strains showed minimal growth in the chemically-defined FMC medium (Terleckyj, Willett, & Shockman, 1975) (Fig. 3A), which does not contain a source of zinc in its original recipe (<1 μM zinc, (Kajfasz et al., 2020). However, supplementation of the FMC medium with as little as 5 μM of ZnSO_4_ restored growth of both *adc* mutants (Fig. 3B). Next, we tested the ability of parent and mutant strains to grow in brain heart infusion (BHI), a complex media containing ~10 μM zinc (Kajfasz et al., 2020) that is routinely used to cultivate *S. mutans* in our laboratory. Based on the behavior of the mutant strains in zinc-supplemented FMC, it was not surprising that both the Δ*adcA* and Δ*adcBC* strains grew well in BHI (Fig. 3C). Next, we tested the ability of the Δ*adcA* and Δ*adcCB* strains to grow in BHI containing purified human calprotectin or in BHI containing the zinc-specific chelator N,N,N’,N’-Tetrakis(2-pyridylmethyl) ethylenediamine (TPEN) (Zhang et al., 2017). In agreement with the role of calprotectin and TPEN in zinc-sequestration, growth of Δ*adcA* and Δ*adcCB* was inhibited by calprotectin (Fig. 2D), or TPEN (Fig. 2E). Finally, the growth defect of the Δ*adcCB* strain under all zinc-depleted conditions were restored in the genetically-complemented Δ*adcCB*^comp^ strain (Fig. 2).

**Figure 3.**
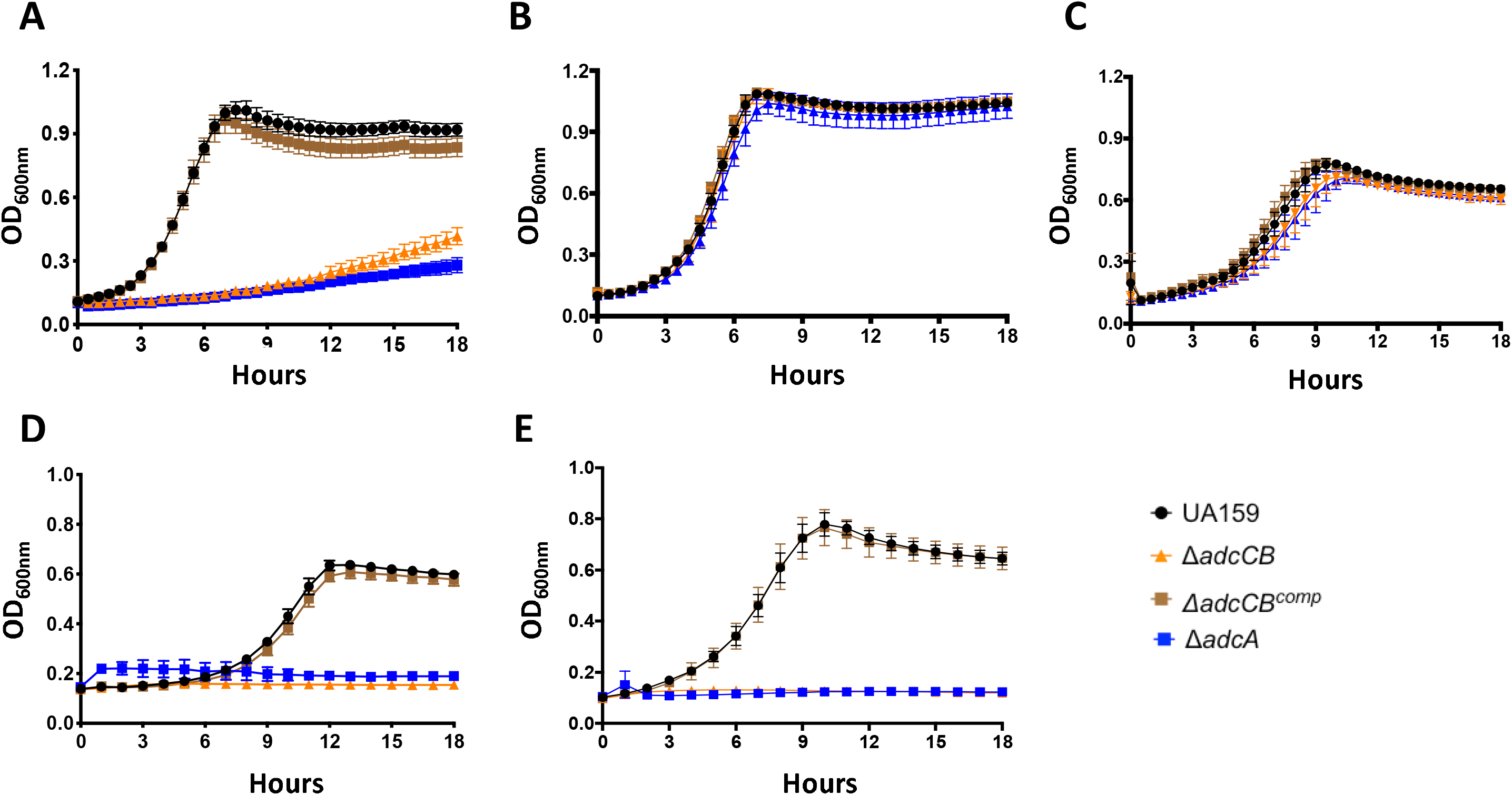
AdcABC mediates the growth of *S. mutans* under zinc-depleted conditions. Growth curves of UA159, Δ*adcA,* Δ*adcCB* or Δ*adcCB*^comp^ in (A) FMC, (B) FMC supplemented with 5 μM ZnSO_4_, (C) BHI, (D) CP/BHI medium containing 200 μg ml^−1^ of human calprotectin, and (E) BHI containing 10 μM TPEN. Curves shown represent average and standard deviation of at least five independent biological replicates.

While AdcAII and Pht-encoding genes are absent in *S. mutans*, a gene (*smu2069*) coding for a hypothetical protein with 37% identity and 60% similarity to the *E.coli* ZupT, a member of the zinc import ZIP family commonly found in gram-negative and in selected gram-positive bacteria (Grass, Wong, Rosen, Smith, & Rensing, 2002; Zackular et al., 2020) was identified through BLAST searches. To probe the possible role of Smu2069 in zinc uptake, we created a strain bearing a *smu2069* deletion in UA159 (Δ*smu2069*) and in the Δ*adcCB* background thereby generating a Δ*adcCB*Δ*smu2069* triple mutant. Inactivation of *smu2069*, alone or in combination with *adcCB*, did not affect zinc uptake as the Δ*smu2069* strain was fully capable to grow in FMC without zinc supplementation and the triple mutant phenocopied the Δ*adcCB* strain (Fig. S2).

Next, we used inductively coupled plasma mass spectrometry (ICP-MS) to determine the intracellular concentration of selected metals (copper, iron, manganese and zinc) in the parent UA159 and derivative strains grown to mid logarithmic phase in BHI or in BHI containing 6 μM TPEN, a concentration permissible to the growth of the Δ*adcCB* and Δ*adcA* strains (data not shown). As compared to UA159, both Δ*adcCB* and Δ*adcA* mutants showed a relatively small, yet significant, reduction in intracellular zinc when grown in plain BHI (Fig. 4A). However, the addition of TPEN to the growth media further increased this difference with both Δ*adcCB* and Δ*adcA* strains showing ~3-fold reduction in intracellular zinc when compared to the parent strain (Fig. 4B). The addition of 6 μM TPEN to the growth media did not affect growth nor the intracellular zinc content of the parent strain confirming that AdcABC alone can maintain intracellular zinc homeostasis under zinc-restricted conditions. Finally, intracellular copper and iron concentrations were not markedly different in parent and mutant strains (data not shown), but there were significant increases in intracellular manganese in Δ*adcCB* (~30% in BHI and ~20% in BHI+TPEN) with the Δ*adcA* strain showing a similar trend in BHI+TPEN (Fig. 4C-D). Collectively, these results indicate that growth of *S. mutans* in zinc-restricted environments is primarily mediated by AdcABC.

**Figure 4.**
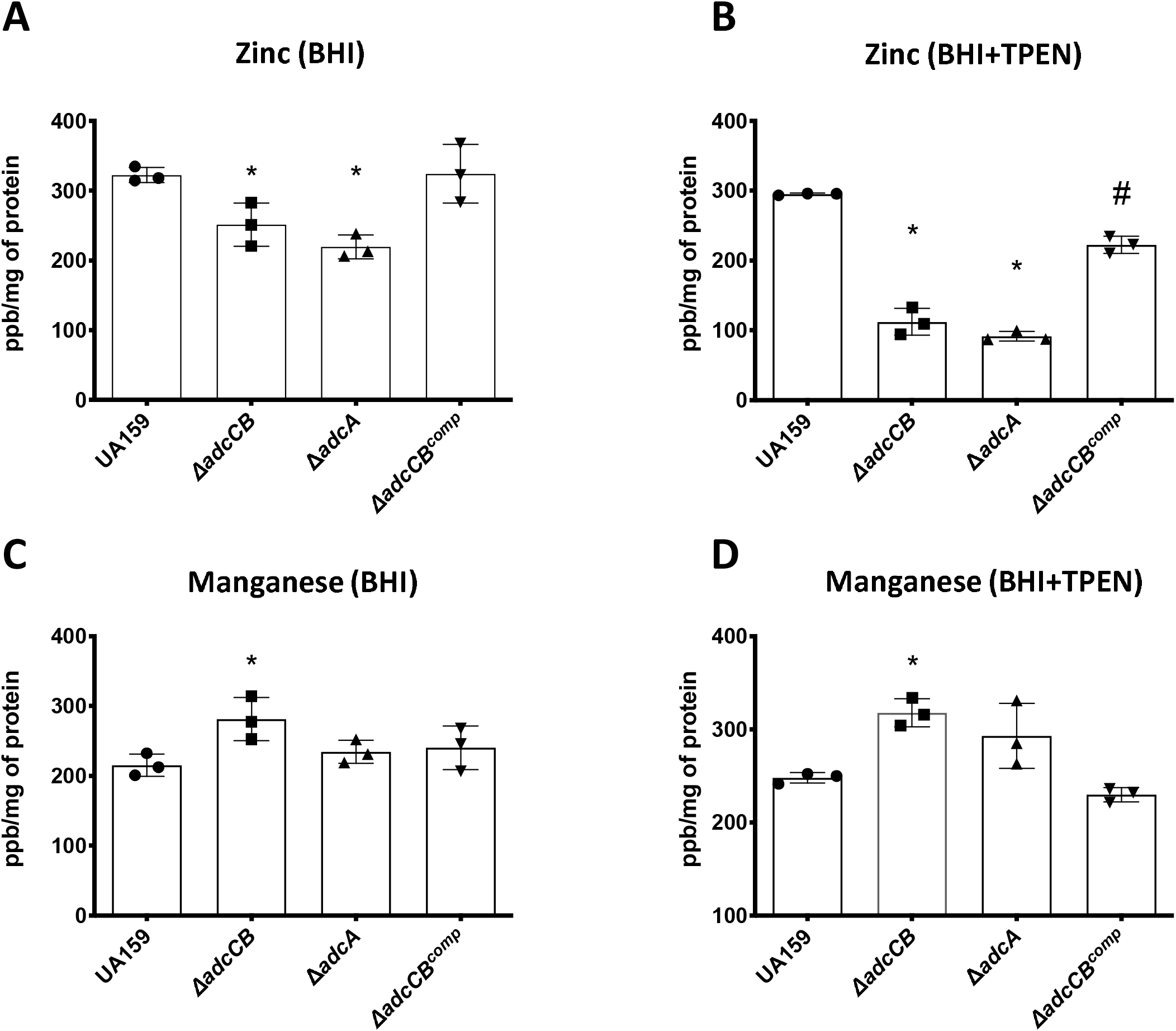
ICP-MS quantifications of intracellular zinc and manganese in the UA159, Δ*adcA,* Δ*adcCB* or Δ*adcCB*^comp^ strains. The bar graphs indicate zinc or manganese levels in cells grown in BHI (A and C) or BHI containing 6 μM TPEN (B and D). Data represent averages and standard deviations of three independent biological replicates. One-way ANOVA was used to compare the metal content of mutants and UA159 (*) and between Δ*adcBC* and Δ*adcCB*^comp^ (#). A *p* value <0.05 was considered significant.

### *The* ΔadcC*B mutant is hypersensitive to high manganese concentrations*

The ICP-MS quantifications indicate that zinc:manganese ratio is drastically different in the Δ*adc* strains when compared to the parent strain, particularly under TPEN-mediated zinc restriction. While zinc:manganese ratios of UA159 and Δ*adc* strains grown in plain BHI was about 1:1 (1:0.7 for UA159 and ~1:1 for Δ*adcA* Δ*ad*c*CB*), there was ~3 times more manganese than zinc (1:3 ratio) in the Δ*adc* strains when grown in BHI+TPEN and a well-balanced 1:1 ratio for UA159. Interestingly, a notable increase in intracellular manganese was also observed in a *S. pneumoniae* double Δ*adcA*Δ*adcAII* strain which, as expected, has a major defect in zinc uptake (Bayle et al., 2011). While manganese is an essential micronutrient to bacteria and lactic acid bacteria are notorious for having a high demand for manganese (Archibald, 1986), recent studies revealed that strains of *S. pneumoniae*, *S. mutans* and *Enterococcus faecalis* lacking the manganese exporter MntE can be intoxicated by manganese (Lam, Wong, Chong, & Kline, 2020; Martin, Lisher, Winkler, & Giedroc, 2017; O’Brien, Pastora, Stoner, & Spatafora, 2020). Using a plate titration assay, we showed that the Δ*adcCB* strain was hypersensitive to high concentrations of manganese (Fig. 5). This hypersensitivity was fully rescued by addition of 20 μM ZnSO_4_ to the growth media directly linking manganese toxicity to a disruption of zinc:manganese balance.

**Figure 5.**
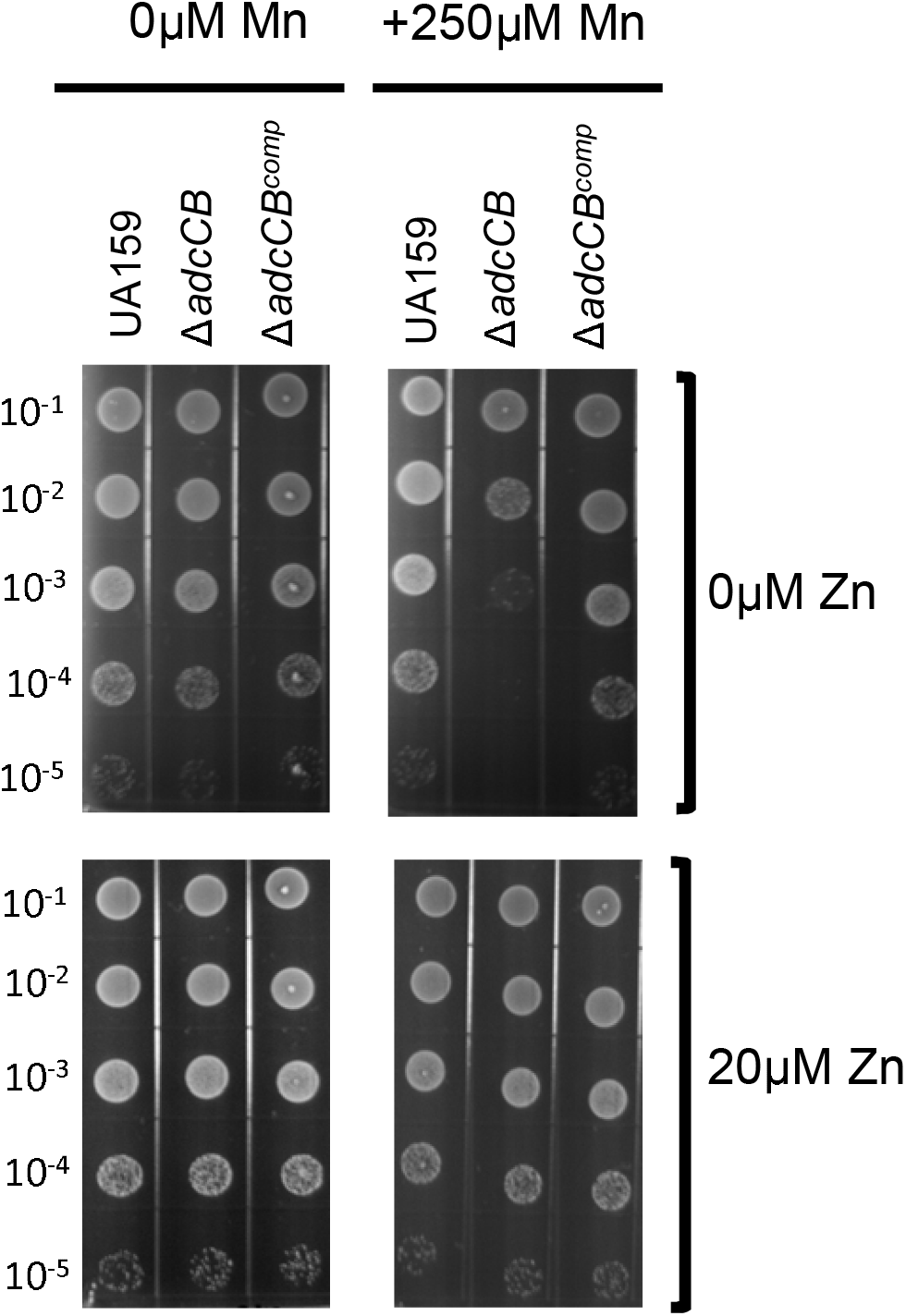
The Δ*adcCB* was hypersensitive to manganese. Mid-log grown cultures of UA159, Δ*adcCB* or Δ*adcCB*^comp^ were serially diluted and spotted on BHI agar containing different concentrations of manganese or zinc as indicated in the figure labels.

### The ΔadcCB strain showed an impaired ability to colonize the rat tooth surface

To determine the significance of zinc transport in oral infection by *S. mutans*, we utilized a rat oral colonization model to test the capacity of the Δ*adcCB* strain to colonize the teeth of rats fed a metal balanced diet (~1.25 nM ZnCO_3_) containing 12% sucrose to facilitate the establishment of *S. mutans* in dental plaque. As compared to the parent strain, the Δ*adcCB* strain was recovered in significantly fewer numbers (~2 logs) two weeks after the first day of infection, with no colonies recovered in two of the eight infected animals (Fig. 6A). Moreover, while *S. mutans*-like colonies accounted for 40 to 60% of the total flora recovered from animals infected with the parent strain, less than 1% of the recovered flora of animals infected with the Δ*adcCB* strain corresponded to *S. mutans*-like colonies (Fig. 6B). All in all, this result indicates that zinc is a growth-limiting factor within the oral biofilm and that the ability to scavenge environmental zinc via the AdcABC transporter is critical to the cariogenic potential of *S. mutans*.

**Figure 6.**
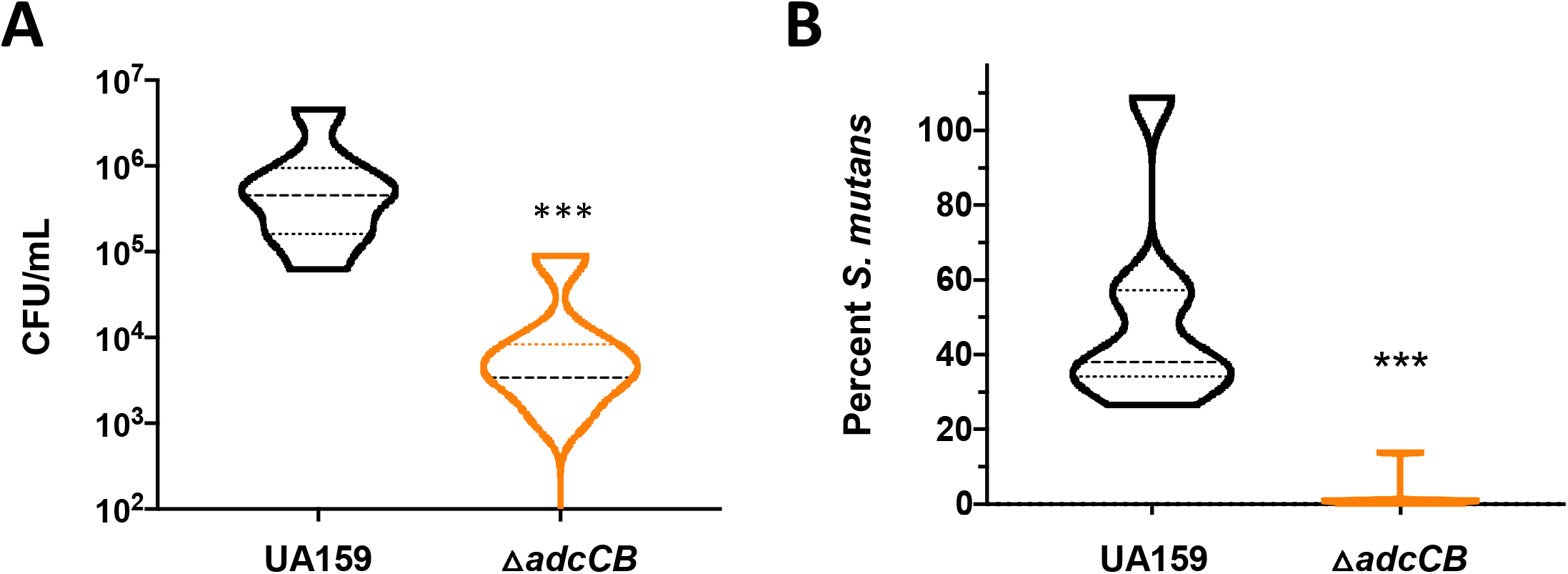
Colonization of *S. mutans* UA159 or Δ*adcBC* on the teeth of rats. (A) Total *S. mutans* colonies recovered from rat jaws by plating on MSB agar, and (B) percentage of *S. mutans* colonies among total recovered flora plated on blood agar plates shown as violin plots. (*) Indicates statistical significance (*p* = 0.0003 (A) and 0.0002 (B) by Mann-Whitney U test).

### *Expression of two virulence-related traits were impaired in the* ΔadcA *and* ΔadcCB *strains*

The ability to cope with environmental stresses, particularly low pH and oxidative stress, and to form robust biofilms in the presence of sucrose are key virulence traits of *S. mutans* (Bowen et al., 2018; Lemos et al., 2019). To investigate whether the oral colonization defect of the Δ*adcCB* strain could be linked to poor expression one or more of these virulence attributes, we first assessed the ability of Δ*adcA* and Δ*adcCB* to tolerate acid or H_2_O_2_ stresses using both growth curve and disc diffusion assays. While the two mutants grew as well as the UA159 parent strain at low pH conditions (data now shown), both showed larger growth inhibition zones when exposed to H_2_O_2_ in a disc diffusion assay (data not shown) and longer lag phase and slightly slower growth rates when grown in FMC supplemented with 5 μM ZnSO_4_, the minimal amount of zinc needed to support growth of the Δ*adc* strains, in the presence of a sub-inhibitory concentration of H_2_O_2_ (Fig. 7A). Notably, increasing the final concentration of ZnSO_4_ from 5 to 20 μM abolished the increased H_2_O_2_ sensitivity of both mutants (Fig. 7B). Next, we tested the ability of the mutants to form biofilm on saliva-coated microtiter plate wells using 1% sucrose as the sugar source. After 24 hours of incubation, both Δ*adcBC* and Δ*adcA* showed a small but significant defect in biofilm formation when grown in FMC containing 5 μM ZnSO_4_ but not 20 μM ZnSO_4_ (Fig. 7C). Collectively, these results serve to further support the oral colonization defect of the Δ*adcBC* strain in the murine model.

**Figure 7.**
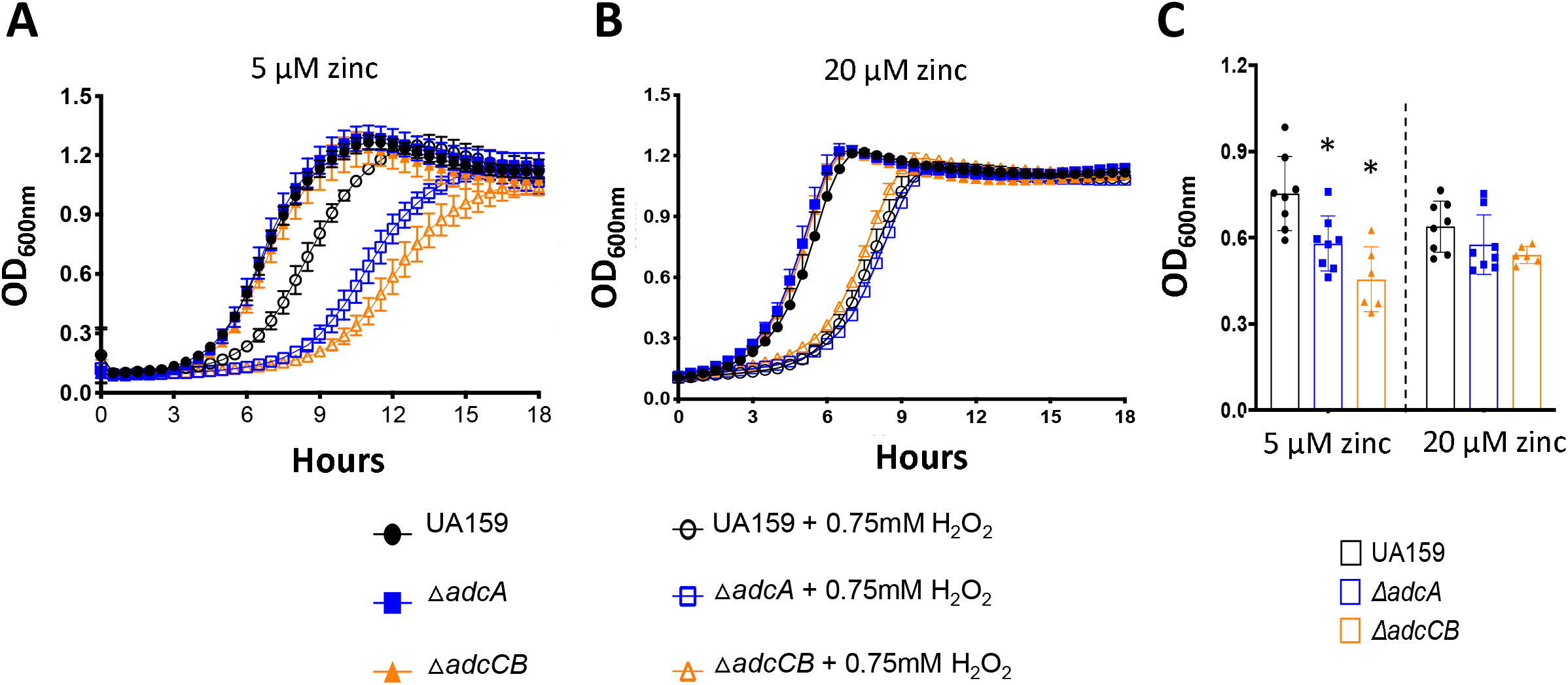
Expression of virulence-related attributes in the UA159, Δ*adcA* and Δ*adcCB* strains. (A) Growth in in the presence of a sub-inhibitory concentration of H_2_O_2_ (0.75 mM) in FMC supplemented with 5 μM (A) or 20 μM ZnSO_4_ (B). Curves shown represent average and standard deviation of at least five independent biological replicates. (C) 24-h biofilms formed on the surface of saliva-coated microtiter plate wells form cells grown in FMC supplemented with 1% sucrose in presence of varying amount of Zn. (*) Indicates statistical significance when compared to UA159 strain (*p* < 0.05, one-way ANOVA).

## Discussion

Despite the critical importance of metals to bacterial virulence, little is known about the mechanisms of metal homeostasis in oral bacteria. In this report, we identified and characterized a high-affinity zinc transporter of *S. mutans*. Inactivation of *adcA* or *adcCB* rendered *S. mutans* unable to grow in the presence of zinc chelators, including calprotectin which can bind zinc in the picomolar range and is produced by host immune cells, specially neutrophils (Brophy, Hayden, & Nolan, 2012; Nakashige et al., 2015; Zackular et al., 2015). Different than other streptococci, the genome of *S. mutans* lacks a second copy of *adcA* or of polyhistidine triad (*pht*) genes. More recently, zinc uptake machineries that resemble siderophore-mediated iron uptake systems were identified in several bacteria including *Staphylococcus aureus* and *Pseudomonas aeruginosa* (Ghssein et al., 2016; Mastropasqua et al., 2017). This system is comprised of small molecules with high zinc affinity (zincophores) that are synthesized in the cytoplasm, exported and re-internalized as a zincophore-zinc complex by a dedicated import system (Grim et al., 2020). Despite evidence that zincophore-mediated uptake systems are produced in a wide array of bacteria, zincophore biosynthetic gene cluster as well as zincophore export and import genes are absent in streptococcal genomes (Morey & Kehl-Fie, 2020). Collectively, our phenotypic characterizations and bioinformatic analysis indicate that the AdcABC transporter is the only high-affinity zinc import system of *S. mutans*.

As the second most abundant trace metals in the human body, zinc is also present in oral fluids and tissues, including the teeth enamel surface, dental plaque and saliva. Independent reports have shown that zinc is detected in the low micromolar range in human saliva and at much higher concentrations in dental plaque, with some studies detecting millimolar levels of zinc in plaque samples (Lynch, 2011). However, the bioavailability of zinc in saliva or in dental plaque is unknown. Our *in vivo* study indicates that zinc is not readily available in dental plaque as the ability of the Δ*adcCB* strain to colonize the teeth of rats fed a zinc-balanced cariogenic diet was severely compromised (Fig. 6). Mammalian hosts utilize a number of mechanisms to restrict zinc access for invading microbes, from zinc tissue/cell reallocation, mediated by two families of zinc transporters, to zinc sequestration/ chelation mechanisms via production of calprotectin (Zackular et al., 2015). While calprotectin can be present in oral fluids and tissues, particularly during inflammatory processes such as periodontitis, oral cancer and oropharyngeal candidiasis (Holmstrom et al., 2019; Sweet, Denbury, & Challacombe, 2001), there is no evidence of increases in salivary calprotectin levels in dental caries. Moreover, recent studies indicated that mildly acidic conditions compromise the ability of calprotectin to chelate manganese but not zinc (Rosen & Nolan, 2020). In addition, salivary glands synthesize metallothioneins, a family of low molecular weight cysteine-rich proteins that scavenge free radicals and can also chelate zinc and copper in tissues (Rahman & Karim, 2018). While the capacity of human metallothionein to restrict bacterial growth through metal sequestration is unclear, salivary metallothionein levels has been shown to increase after dental pulp injury in rats (Izumi, Eida, Matsumoto, & Inoue, 2007). To obtain preliminary insight into the host-derived zinc sequestration mechanisms in the oral cavity, we took advantage of the availability of stored saliva samples from caries-free and caries active subjects that had been previously collected for a recently completed clinical study (Garcia *et al*, manuscript in preparation), and used ELISAs to measure the levels of calprotectin and metallothionein in these samples. While calprotectin was below detection levels in most samples, there was measurable quantities of metallothionenin in the saliva samples from both subject groups (Fig. S3). However, the differences in the levels of metallothionenin between the two groups were not significant suggesting that metallothionein levels do not increase in response to caries. Given the very low levels of calprotectin that we found in saliva and recent observations that the manganese-binding affinity of calprotectin is compromised at low pH (Rosen & Nolan, 2020), we speculate that zinc restriction in the oral cavity is primarily mediated by metallothioneins.

In addition to the host-driven mechanisms of metal sequestration, bacteria in polymicrobial biofilms such as dental plaque must also compete with the other oral residents for nutrients. Interestingly, a recent study showed that transcription of the *S. mutans adcA* and *adcB* genes was induced (about 6- and 3-fold, respectively) when *S. mutans* was co-cultured with *Streptococcus* A12, a health-associated peroxigenic oral streptococci (Kaspar, Lee, Richard, Walker, & Burne, 2020). It should be noted that, in principle, peroxigenic streptococci are better equipped to scavenge zinc than *S. mutans* due to the concerted effort of AdcAII and Pht proteins in addition to the AdcABC system. Another important aspect to consider when it comes to zinc availability is the effect of pH on trace metal solubility. At acidic pH values (pH<6), most free zinc found in saliva is in the aquated Zn^2+^ form, but it sharply decreases if the environmental pH increases above 6 (Rahman, Hossain, Pin, & Yahya, 2019). Thus, it is conceivable that as the plaque pH lowers as a result of bacterial metabolism, zinc availability to *mutans* might increase due to increased solubilization and concomitant reduction of acid-sensitive competitors. In addition, it is conceivable that net H_2_O_2_ production in dental plaque due to the presence of peroxigenic oral bacteria will stimulate metallothionein synthesis by salivary glands thereby limiting the availability of free zinc in the oral cavity. Studies to determine how host-pathogen and microbe-microbe interactions influence the ability of *S. mutans* to maintain zinc homeostasis in dental plaque will soon be underway.

While this study focused on the bacterial zinc acquisition mechanisms, it should be noted that, similar to other trace metals, excess zinc is poisonous to microbes. In addition to being essential to all forms of life, zinc is also recognized for having antimicrobial properties and to stimulate immune cell function such that zinc-containing products have been used in wound healing, as an adjuvant for the treatment of the common cold, and as antimicrobials. In the context of oral health, zinc has been incorporated to mouthwash and toothpaste formulations to assist in the control of calculus formation, gingivitis and halitosis, while the role of zinc as an anticaries agent is controversial (Barnes, Richter, Bastin, Lambert, & Xu, 2008; Bates & Navia, 1979; Giertsen, 2004; Harrap, Best, & Saxton, 1984; Li et al., 2015). Few studies have probed the consequences of high zinc concentrations to the physiology of oral streptococci, with *in vitro* studies showing that high zinc concentrations inhibit their metabolism (He, Pearce, & Sissons, 2002; Phan, Buckner, Sheng, Baldeck, & Marquis, 2004). Because the incorporation of zinc to routinely used oral health care products is predicted to disturb dental plaque ecology, future studies to determine the importance of environmental zinc in the caries process should also explore zinc excess conditions. From a translational standpoint, approaches to restrict the access of oral bacteria to zinc or, conversely, intoxicate them with zinc may prove effective in the control of dental caries.

## Experimental procedures

### Bacterial strains growth conditions

The *S. mutans* strains used in this study are listed in Table 1. Strains were routinely grown in brain heart infusion (BHI) medium at 37°C in a 5% CO_2_ incubator. When appropriate, spectinomycin (1 mg ml^−1^), kanamycin (1 mg ml^−1^) or erythromycin (10 μg ml^−1^) was added to the growth media. In addition, the chemically defined FMC medium (Terleckyj et al., 1975) was used to determine the minimal amounts of zinc that support growth of the mutant strains. Growth curves under different stress conditions and using media (BHI or FMC) with different amounts of zinc were obtained using an automated growth reader set to 37°C (Bioscreen C; Oy Growth Curves Ab, Ltd.) as described previously (Kajfasz et al., 2010). Briefly, overnight cultures grown in BHI were diluted 1:50 in to the appropriate medium in the wells of a microtiter plate with an overlay of sterile mineral oil added to each well to minimize oxidative stress. Growth in the presence of calprotectin requires the use of 38% bacterial medium and 62% CP buffer (20 mM Tris [pH 7.5], 100 mM NaCl, 3 mM CaCl_2_, 5 mM β-mercaptoethanol). To promote growth of *S. mutans* in the CP medium, 3X concentrated BHI medium was used in combination with the CP buffer. To determine the sensitivity of the different strains to manganese, cultures were grown in BHI to an OD_600_ of 0.5, serially diluted and 10 μl of each dilution spotted onto BHI agar supplemented with 250 μM MnSO_4_ with or without additional zinc (20 μM ZnSO_4_). Plates were incubated for 24 hours at 37°C in a 5% CO_2_ incubator before they were photographed.

**Table 1.**
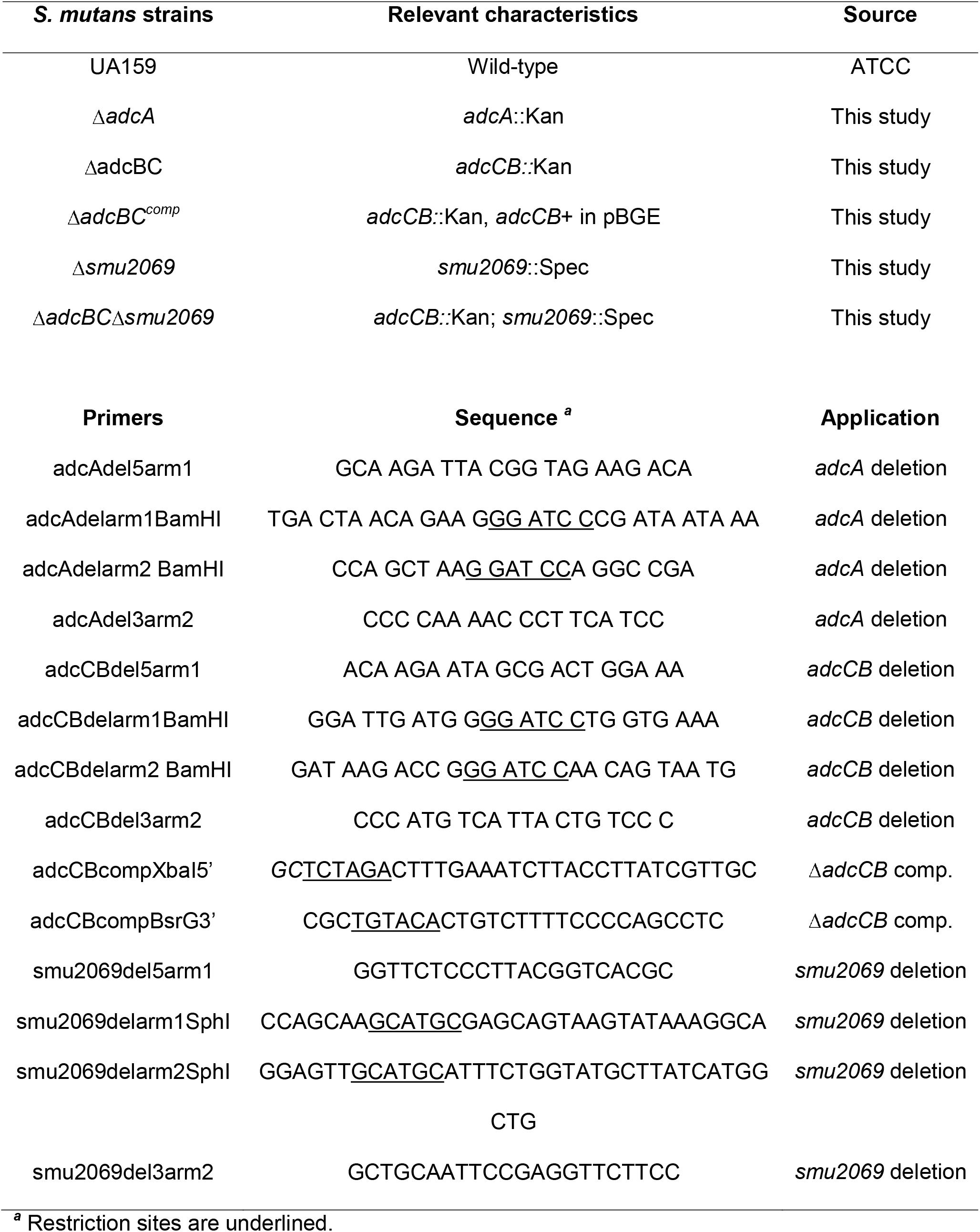
Bacterial strains and primers used in the study.

### Construction of mutant strains

Deletion strains lacking the *adcA*, *adcCB* or *smu2069* genes were obtained using a PCR ligation mutagenesis approach (Lau, Sung, Lee, Morrison, & Cvitkovitch, 2002). Briefly, PCR fragments flanking the region to be deleted were ligated to a nonpolar kanamycin (*adcA* and *adcCB*) or spectinomycin (*smu2069*) resistance cassette and the ligation mixture used to transform *S. mutans* UA159 according to an established protocol (Lau et al., 2002). Mutant strains were selected on BHI agar supplemented with the appropriate antibiotic and confirmed by PCR analysis and DNA sequencing. To generate complemented strains, the full length *adcRCB* operon was amplified by PCR and cloned into the integration vector pBGE (Zeng & Burne, 2009) to yield plasmid pBGE-*adcRCB*. The plasmid was propagated in *E. coli* DH10B and used to transform the *S. mutans* Δ*adcCB* strain for integration at the *gtfA* locus. All primers used in this study are listed in Table 1.

### ICP-MS analysis

The bacterial intracellular metal content was determined using Inductively Coupled Plasma Mass Spectrometry (ICP-MS). Briefly, cultures were grown in BHI or BHI containing 6 μM TPEN to an OD_600_ of 0.4, harvested by centrifugation at 4°C for 15□min at 4,000 rpm, washed in phosphate-buffered saline (PBS) supplemented with 0.2□mM EDTA to chelate extracellular divalent cations followed by a wash in PBS alone. The cell pellets were resuspended in HNO_3_ and metal composition was quantified using an Agilent 7900 ICP mass spectrometer. Metal concentrations were then normalized to total protein content as determined by the bicinchoninic acid (BCA) assay (Pierce).

### Biofilm assay

The ability of the *S. mutans* strains to form biofilms on saliva-coated wells of polystyrene microtiter plates was assessed as described elsewhere (Kajfasz et al., 2020). The wells were first coated with 100□μl of sterile clarified and pooled human saliva (IRB# 201600877) for 30 min at room temperature. Strains were grown in BHI to an OD_600_ of 0.5 and diluted 1:100 in FMC containing 1% sucrose (FMC-S) and supplemented with 5 or 20 μM ZnSO_4_, and 200 μl of each bacterial suspension added to the saliva-coated wells. After incubation at 37°C in a 5% CO_2_ incubator for 24 hours, wells were washed twice with sterile water to remove planktonic and loosely bound bacteria, and adherent (biofilm) cells stained with 0.1% crystal violet for 15□min. The bound dye was eluted in a 33% acetic acid solution, and the total biofilm estimated by measuring the optical density of the dissolved dye at 575□nm.

### Rat tooth colonization assay

The ability of the mutant strains to colonize the teeth of rats was evaluated using an established model of dental caries (Miller et al., 2015) with some modifications. Briefly, specific pathogen-free Sprague-Dawley rat pups were purchased with their dams from Jackson Laboratories and screened for the presence of mutans streptococci by plating on mitis salivarius (MS) agar upon arrival. Prior to infection, pups and dams received 0.8 mg ml^−1^ sulfamethoxazole and 0.16 mg ml^−1^ trimethoprim in the drinking water for 3 days to suppress endogenous flora and facilitate infection. Pups aged 15 days and dams were taken off antibiotics on day 4 randomly placed into experimental groups, and infected for four consecutive days by means of cotton swab saturated with actively growing *S. mutans* UA159 or Δ*adcBC* cultures. During the days of infection, animals were fed a 12% sucrose powdered diet (ENVIGO diet, catalog # TD.190707) and 5% (wt/vol) sterile sucrose-water *ad libitum*, and the 12% sucrose diet with sterile water in the days after infection. The experiment proceeded for 10 additional days, at the end of which the animals were euthanized by CO_2_ asphyxiation, and the lower jaws surgically removed for microbiological assessment. Jaw sonicates were subjected to 10-fold serial dilution in PBS and plated on MS agar to determine *S. mutans* counts. The number of *S. mutans* recovered from the animals was expressed as CFU ml-1 of jaw sonicate. This study was reviewed and approved by the University of Florida Institutional Animal Care and Use Committee (protocol # 201810421).

### Statistical analysis

Data were analyzed using GraphPad Prism 9.0 software unless otherwise stated. Differences in intracellular metal content and biofilm formation were determined via ordinary one-way ANOVA. The rat colonization study was subjected to the Mann-Whitney U test. In all cases, *p* values < 0.05 were considered significant.

## Acknowledgements

Purified calprotectin was provided by Walter Chazin at Vanderbilt University. This study was supported by NIH-NIDCR award DE019783 to J.A.L.

## Author contributions

TG, JKK, JA and JAL designed the study; TG, AMP, JKK and JA performed the experiments, TG, JKK, JA and JAL wrote the manuscript.

---

**Figure S1.**
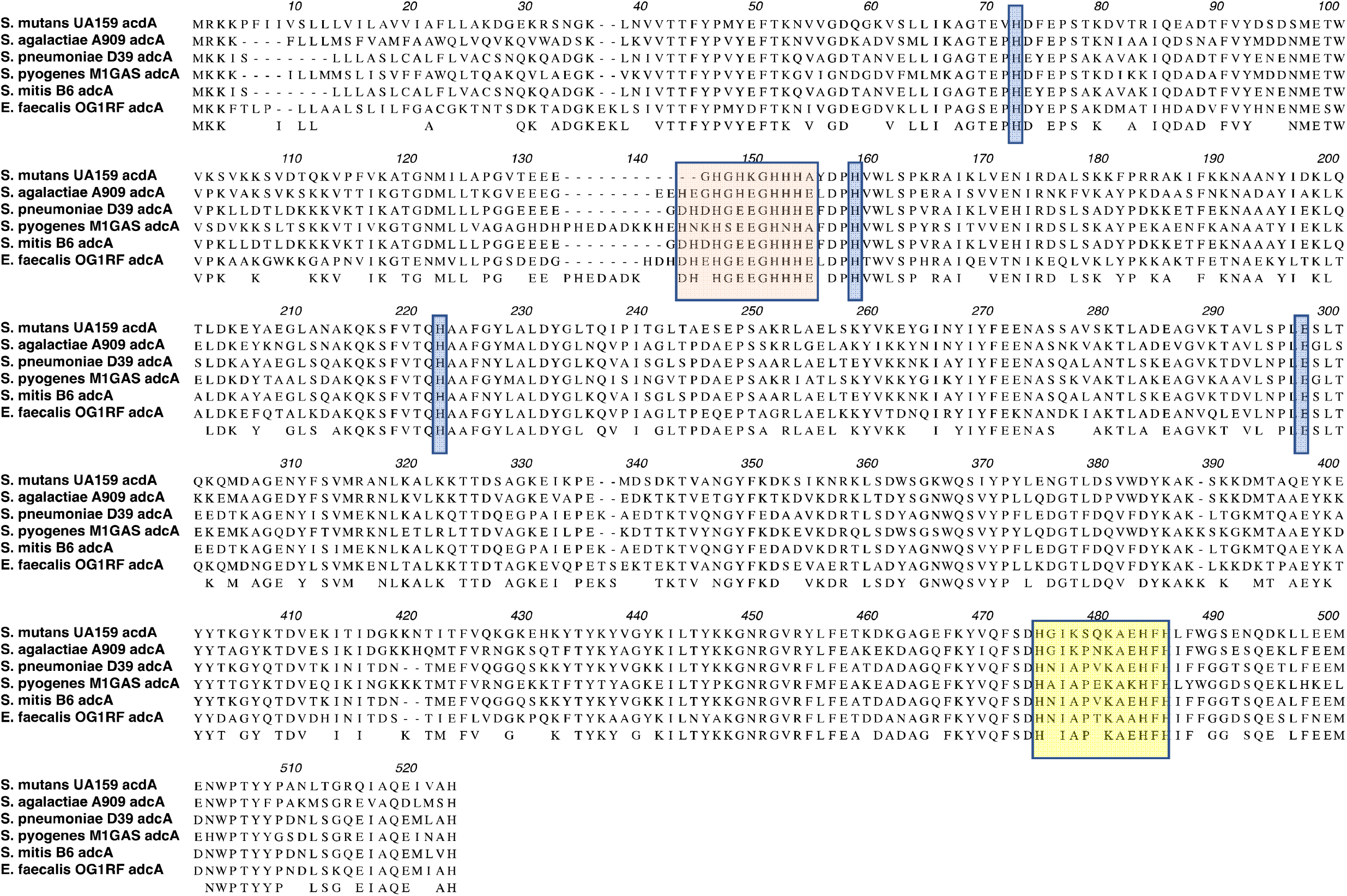
Amino acid alignment of several AdcA proteins from several streptococci and *E. faecalis*. Orange shaded residues are the N-terminal histidine rich metal-binding motif, yellow boxed residues depict the C-terminal ZinT domain, the glutamic acid and additional histidine residues that aid in metal recruitment are indicated in blue shades.

**Figure S2.**
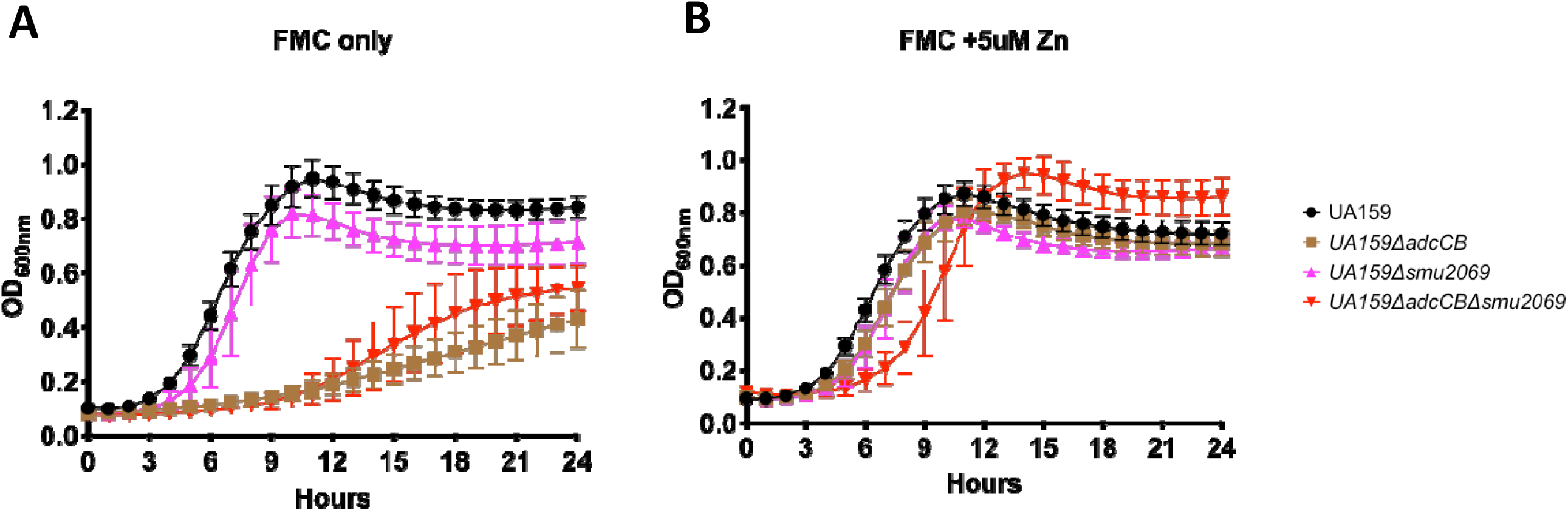
Growth of the *S. mutans* UA159, Δ*adcCB*, Δ*smu2069,* or Δ*adcBC*Δ*smu2069* strains in (A) FMC or (B) FMC supplemented with 5 μM zinc.

**Figure S3.**
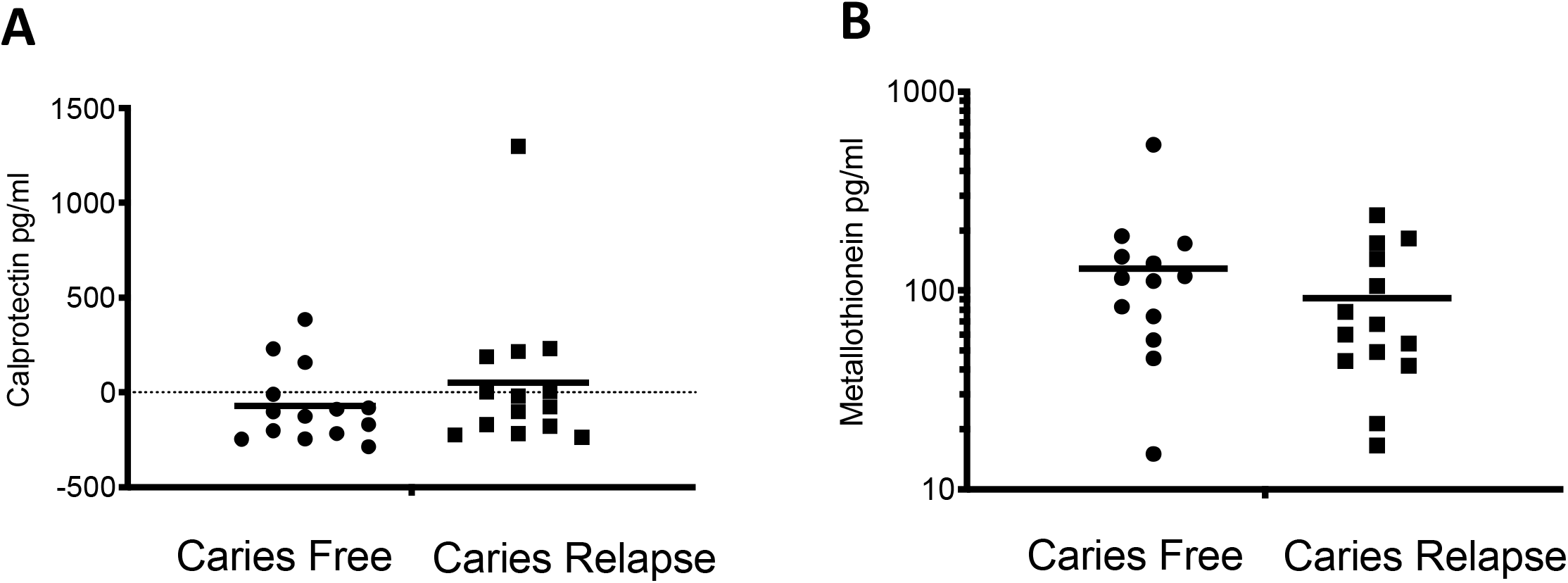
Salivary levels of calprotectin (A) and metallothionein (B) in human saliva determined by ELISA.

## References

Archibald, F. (1986). Manganese: its acquisition by and function in the lactic acid bacteria. Crit Rev Microbiol, 13(1), 63–109. doi:10.3109/10408418609108735

Barnes, V. M., Richter, R., Bastin, D., Lambert, P., & Xu, T. (2008). Dental plaque control effect of a zinc citrate dentifrice. J Clin Dent, 19(4), 127–130.

Bates, D. G., & Navia, J. M. (1979). Chemotherapeutic effect of zinc on streptococcus mutans and rat dental caries. Arch Oral Biol, 24(10-11), 799–805. doi:10.1016/0003-9969(79)90041-4

Bayle, L., Chimalapati, S., Schoehn, G., Brown, J., Vernet, T., & Durmort, C. (2011). Zinc uptake by Streptococcus pneumoniae depends on both AdcA and AdcAII and is essential for normal bacterial morphology and virulence. Mol Microbiol, 82(4), 904–916. doi:10.1111/j.1365-2958.2011.07862.x

Bersch, B., Bougault, C., Roux, L., Favier, A., Vernet, T., & Durmort, C. (2013). New insights into histidine triad proteins: solution structure of a Streptococcus pneumoniae PhtD domain and zinc transfer to AdcAII. PLoS One, 8(11), e81168 . doi:10.1371/journal.pone.0081168

Bowen, W. H., Burne, R. A., Wu, H., & Koo, H. (2018). Oral Biofilms: Pathogens, Matrix, and Polymicrobial Interactions in Microenvironments. Trends Microbiol, 26(3), 229–242. doi:10.1016/j.tim.2017.09.008

Brophy, M. B., Hayden, J. A., & Nolan, E. M. (2012). Calcium ion gradients modulate the zinc affinity and antibacterial activity of human calprotectin. J Am Chem Soc, 134(43), 18089–18100. doi:10.1021/ja307974e

Brown, L. R., Gunnell, S. M., Cassella, A. N., Keller, L. E., Scherkenbach, L. A., Mann, B., … Thornton, J. A. (2016). AdcAII of Streptococcus pneumoniae Affects Pneumococcal Invasiveness. PLoS One, 11(1), e0146785. doi:10.1371/journal.pone.0146785

Burcham, L. R., Le Breton, Y., Radin, J. N., Spencer, B. L., Deng, L., Hiron, A., … Doran, K. S. (2020). Identification of Zinc-Dependent Mechanisms Used by Group B Streptococcus To Overcome Calprotectin-Mediated Stress. mBio, 11(6). doi:10.1128/mBio.02302-20

Cao, K., Li, N., Wang, H., Cao, X., He, J., Zhang, B., … Sun, X. (2018). Two zinc-binding domains in the transporter AdcA from Streptococcus pyogenes facilitate high-affinity binding and fast transport of zinc. J Biol Chem, 293(16), 6075–6089. doi:10.1074/jbc.M117.818997

Chandrangsu, P., & Helmann, J. D. (2016). Intracellular Zn(II) Intoxication Leads to Dysregulation of the PerR Regulon Resulting in Heme Toxicity in Bacillus subtilis. PLoS Genet, 12(12), e1006515. doi:10.1371/journal.pgen.1006515

Fischer, F., Robbe-Saule, M., Turlin, E., Mancuso, F., Michel, V., Richaud, P., … Vinella, D. (2016). Characterization in Helicobacter pylori of a Nickel Transporter Essential for Colonization That Was Acquired during Evolution by Gastric Helicobacter Species. PLoS Pathog, 12(12), e1006018. doi:10.1371/journal.ppat.1006018

Garcia, E. C., Brumbaugh, A. R., & Mobley, H. L. (2011). Redundancy and specificity of Escherichia coli iron acquisition systems during urinary tract infection. Infect Immun, 79(3), 1225–1235. doi:10.1128/IAI.01222-10

Ghssein, G., Brutesco, C., Ouerdane, L., Fojcik, C., Izaute, A., Wang, S., … Arnoux, P. (2016). Biosynthesis of a broad-spectrum nicotianamine-like metallophore in Staphylococcus aureus. Science, 352(6289), 1105–1109. doi:10.1126/science.aaf1018

Giertsen, E. (2004). Effects of mouthrinses with triclosan, zinc ions, copolymer, and sodium lauryl sulphate combined with fluoride on acid formation by dental plaque in vivo. Caries Res, 38(5), 430–435. doi:10.1159/000079623

Grass, G., Wong, M. D., Rosen, B. P., Smith, R. L., & Rensing, C. (2002). ZupT is a Zn(II) uptake system in Escherichia coli. J Bacteriol, 184(3), 864–866. doi:10.1128/jb.184.3.864-866.2002

Grim, K. P., Radin, J. N., Solorzano, P. K. P., Morey, J. R., Frye, K. A., Ganio, K., … Kehl-Fie, E. (2020). Intracellular Accumulation of Staphylopine Can Sensitize Staphylococcus aureus to Host-Imposed Zinc Starvation by Chelation-Independent Toxicity. J Bacteriol, 202(9). doi:10.1128/JB.00014-20

Harrap, G. J., Best, J. S., & Saxton, C. A. (1984). Human oral retention of zinc from mouthwashes containing zinc salts and its relevance to dental plaque control. Arch Oral Biol, 29(2), 87–91. doi:10.1016/0003-9969(84)90110-9

He, G., Pearce, E. I., & Sissons, C. H. (2002). Inhibitory effect of ZnCl(2) on glycolysis in human oral microbes. Arch Oral Biol, 47(2), 117–129. doi:10.1016/s0003-9969(01)00093-0

Holmstrom, S. B., Lira-Junior, R., Zwicker, S., Majster, M., Gustafsson, A., Akerman, S., … Bostrom, E. A. (2019). MMP-12 and S100s in saliva reflect different aspects of periodontal inflammation. Cytokine, 113, 155–161. doi:10.1016/j.cyto.2018.06.036

Imlay, J. A. (2014). The mismetallation of enzymes during oxidative stress. J Biol Chem, 289(41), 28121–28128. doi:10.1074/jbc.R114.588814

Izumi, T., Eida, T., Matsumoto, N., & Inoue, H. (2007). Immunohistochemical localization of metallothionein in dental pulp after cavity preparation of rat molars. Oral Surg Oral Med Oral Pathol Oral Radiol Endod, 104(4), e133–137. doi:10.1016/j.tripleo.2007.04.023

Juttukonda, L. J., & Skaar, E. P. (2015). Manganese homeostasis and utilization in pathogenic bacteria. Mol Microbiol, 97(2), 216–228. doi:10.1111/mmi.13034

Kajfasz, J. K., Katrak, C., Ganguly, T., Vargas, J., Wright, L., Peters, Z. T., … Lemos, J. A. (2020). Manganese Uptake, Mediated by SloABC and MntH, Is Essential for the Fitness of Streptococcus mutans. mSphere, 5(1). doi:10.1128/mSphere.00764-19

Kajfasz, J. K., Rivera-Ramos, I., Abranches, J., Martinez, A. R., Rosalen, P. L., Derr, A. M., … Lemos, J. A. (2010). Two Spx proteins modulate stress tolerance, survival, and virulence in Streptococcus mutans. J Bacteriol, 192(10), 2546–2556. doi:10.1128/JB.00028-10

Kaspar, J. R., Lee, K., Richard, B., Walker, A. R., & Burne, R. A. (2020). Direct interactions with commensal streptococci modify intercellular communication behaviors of Streptococcus mutans. ISME J. doi:10.1038/s41396-020-00789-7

Kehl-Fie, T. E., & Skaar, E. P. (2010). Nutritional immunity beyond iron: a role for manganese and zinc. Curr Opin Chem Biol, 14(2), 218–224. doi:10.1016/j.cbpa.2009.11.008

Koh, E. I., Hung, C. S., Parker, K. S., Crowley, J. R., Giblin, D. E., & Henderson, J. P. (2015). Metal selectivity by the virulence-associated yersiniabactin metallophore system. Metallomics, 7(6), 1011–1022. doi:10.1039/c4mt00341a

Lam, L. N., Wong, J. J., Chong, K. K. L., & Kline, K. A. (2020). Enterococcus faecalis Manganese Exporter MntE Alleviates Manganese Toxicity and Is Required for Mouse Gastrointestinal Colonization. Infect Immun, 88(6). doi:10.1128/IAI.00058-20

Lau, P. C., Sung, C. K., Lee, J. H., Morrison, D. A., & Cvitkovitch, D. G. (2002). PCR ligation mutagenesis in transformable streptococci: application and efficiency. J Microbiol Methods, 49(2), 193–205. doi:10.1016/s0167-7012(01)00369-4

Lemos, J. A., Palmer, S. R., Zeng, L., Wen, Z. T., Kajfasz, J. K., Freires, I. A., … Brady, L. J. (2019). The Biology of Streptococcus mutans. Microbiol Spectr, 7(1). doi:10.1128/microbiolspec.GPP3-0051-2018

Li, X., Zhong, Y., Jiang, X., Hu, D., Mateo, L. R., Morrison, B. M., Jr., & Zhang, Y. P. (2015). Randomized clinical trial of the efficacy of dentifrices containing 1.5% arginine, an insoluble calcium compound and 1450 ppm fluoride over two years. J Clin Dent, 26(1), 7–12.

Loisel, E., Chimalapati, S., Bougault, C., Imberty, A., Gallet, B., Di Guilmi, A. M., … Durmort, C. (2011). Biochemical characterization of the histidine triad protein PhtD as a cell surface zinc-binding protein of pneumococcus. Biochemistry, 50(17), 3551–3558. doi:10.1021/bi200012f

Loisel, E., Jacquamet, L., Serre, L., Bauvois, C., Ferrer, J. L., Vernet, T., … Durmort, C. (2008). AdcAII, a new pneumococcal Zn-binding protein homologous with ABC transporters: biochemical and structural analysis. J Mol Biol, 381(3), 594–606. doi:10.1016/j.jmb.2008.05.068

Lonergan, Z. R., & Skaar, E. P. (2019). Nutrient Zinc at the Host-Pathogen Interface. Trends Biochem Sci, 44(12), 1041–1056. doi:10.1016/j.tibs.2019.06.010

Loo, C. Y., Mitrakul, K., Voss, I. B., Hughes, C. V., & Ganeshkumar, N. (2003). Involvement of the adc operon and manganese homeostasis in Streptococcus gordonii biofilm formation. J Bacteriol, 185(9), 2887–2900. doi:10.1128/jb.185.9.2887-2900.2003

Lynch, R. J. (2011). Zinc in the mouth, its interactions with dental enamel and possible effects on caries; a review of the literature. Int Dent J, 61 Suppl 3, 46–54. doi:10.1111/j.1875-595X.2011.00049.x

Martin, J. E., Lisher, J. P., Winkler, M. E., & Giedroc, D. P. (2017). Perturbation of manganese metabolism disrupts cell division in Streptococcus pneumoniae. Mol Microbiol, 104(2), 334–348. doi:10.1111/mmi.13630

Mastropasqua, M. C., D’Orazio, M., Cerasi, M., Pacello, F., Gismondi, A., Canini, A., … Battistoni, A. (2017). Growth of Pseudomonas aeruginosa in zinc poor environments is promoted by a nicotianamine-related metallophore. Mol Microbiol, 106(4), 543–561. doi:10.1111/mmi.13834

Miller, J. H., Aviles-Reyes, A., Scott-Anne, K., Gregoire, S., Watson, G. E., Sampson, E., … Abranches, J. (2015). The collagen binding protein Cnm contributes to oral colonization and cariogenicity of Streptococcus mutans OMZ175. Infect Immun, 83(5), 2001–2010. doi:10.1128/IAI.03022-14

Morey, J. R., & Kehl-Fie, T. E. (2020). Bioinformatic Mapping of Opine-Like Zincophore Biosynthesis in Bacteria. mSystems, 5(4). doi:10.1128/mSystems.00554-20

Moulin, P., Patron, K., Cano, C., Zorgani, M. A., Camiade, E., Borezee-Durant, E., … Hiron, A. (2016). The Adc/Lmb System Mediates Zinc Acquisition in Streptococcus agalactiae and Contributes to Bacterial Growth and Survival. J Bacteriol, 198(24), 3265–3277. doi:10.1128/JB.00614-16

Nakashige, T. G., Zhang, B., Krebs, C., & Nolan, E. M. (2015). Human calprotectin is an iron-sequestering host-defense protein. Nat Chem Biol, 11(10), 765–771. doi:10.1038/nchembio.1891

O’Brien, J., Pastora, A., Stoner, A., & Spatafora, G. (2020). The S. mutans mntE gene encodes a manganese efflux transporter. Mol Oral Microbiol, 35(3), 129–140. doi:10.1111/omi.12286

Ong, C. Y., Berking, O., Walker, M. J., & McEwan, A. G. (2018). New Insights into the Role of Zinc Acquisition and Zinc Tolerance in Group A Streptococcal Infection. Infect Immun, 86(6). doi:10.1128/IAI.00048-18

Palmer, L. D., & Skaar, E. P. (2016). Transition Metals and Virulence in Bacteria. Annu Rev Genet, 50, 67–91. doi:10.1146/annurev-genet-120215-035146

Pant, S., Patel, N. J., Deshmukh, A., Golwala, H., Patel, N., Badheka, A., … Mehta, J. L. (2015). Trends in infective endocarditis incidence, microbiology, and valve replacement in the United States from 2000 to 2011. J Am Coll Cardiol, 65(19), 2070–2076. doi:10.1016/j.jacc.2015.03.518

Phan, T. N., Buckner, T., Sheng, J., Baldeck, J. D., & Marquis, R. E. (2004). Physiologic actions of zinc related to inhibition of acid and alkali production by oral streptococci in suspensions and biofilms. Oral Microbiol Immunol, 19(1), 31–38. doi:10.1046/j.0902-0055.2003.00109.x

Rahman, M. T., Hossain, A., Pin, C. H., & Yahya, N. A. (2019). Zinc and Metallothionein in the Development and Progression of Dental Caries. Biol Trace Elem Res, 187(1), 51–58. doi:10.1007/s12011-018-1369-z

Rahman, M. T., & Karim, M. M. (2018). Metallothionein: a Potential Link in the Regulation of Zinc in Nutritional Immunity. Biol Trace Elem Res, 182(1), 1–13. doi:10.1007/s12011-017-1061-8

Reyes-Caballero, H., Guerra, A. J., Jacobsen, F. E., Kazmierczak, K. M., Cowart, D., Koppolu, U. M., … Giedroc, D. P. (2010). The metalloregulatory zinc site in Streptococcus pneumoniae AdcR, a zinc-activated MarR family repressor. J Mol Biol, 403(2), 197–216. doi:10.1016/j.jmb.2010.08.030

Rosen, T., & Nolan, E. M. (2020). Metal Sequestration and Antimicrobial Activity of Human Calprotectin Are pH-Dependent. Biochemistry, 59(26), 2468–2478. doi:10.1021/acs.biochem.0c00359

Sheldon, J. R., & Skaar, E. P. (2019). Metals as phagocyte antimicrobial effectors. Curr Opin Immunol, 60, 1–9. doi:10.1016/j.coi.2019.04.002

Subramanian Vignesh, K., & Deepe, G. S., Jr. (2016). Immunological orchestration of zinc homeostasis: The battle between host mechanisms and pathogen defenses. Arch Biochem Biophys, 611, 66–78. doi:10.1016/j.abb.2016.02.020

Sweet, S. P., Denbury, A. N., & Challacombe, S. J. (2001). Salivary calprotectin levels are raised in patients with oral candidiasis or Sjogren’s syndrome but decreased by HIV infection. Oral Microbiol Immunol, 16(2), 119–123. doi:10.1034/j.1399-302x.2001.016002119.x

Terleckyj, B., Willett, N. P., & Shockman, G. D. (1975). Growth of several cariogenic strains of oral streptococci in a chemically defined medium. Infect Immun, 11(4), 649–655. doi:10.1128/IAI.11.4.649-655.1975

Zackular, J. P., Chazin, W. J., & Skaar, E. P. (2015). Nutritional Immunity: S100 Proteins at the Host-Pathogen Interface. J Biol Chem, 290(31), 18991–18998. doi:10.1074/jbc.R115.645085

Zackular, J. P., Knippel, R. J., Lopez, C. A., Beavers, W. N., Maxwell, C. N., Chazin, W. J., & Skaar, E. P. (2020). ZupT Facilitates Clostridioides difficile Resistance to Host-Mediated Nutritional Immunity. mSphere, 5(2). doi:10.1128/mSphere.00061-20

Zeng, L., & Burne, R. A. (2009). Transcriptional regulation of the cellobiose operon of Streptococcus mutans. J Bacteriol, 191(7), 2153–2162. doi:10.1128/JB.01641-08

Zhang, F., Ma, X. L., Wang, Y. X., He, C. C., Tian, K., Wang, H. G., … Liu, Y. Q. (2017). TPEN, a Specific Zn(2+) Chelator, Inhibits Sodium Dithionite and Glucose Deprivation (SDGD)-Induced Neuronal Death by Modulating Apoptosis, Glutamate Signaling, and Voltage-Gated K(+) and Na(+) Channels. Cell Mol Neurobiol, 37(2), 235–250. doi:10.1007/s10571-016-0364-1

Zheng, B., Zhang, Q., Gao, J., Han, H., Li, M., Zhang, J., … Gao, G. F. (2011). Insight into the interaction of metal ions with TroA from Streptococcus suis. PLoS One, 6(5), e19510. doi:10.1371/journal.pone.0019510

